# iREAD: A Tool for Intron Retention Detection from RNA-seq Data

**DOI:** 10.1101/135624

**Authors:** Hong-Dong Li, Cory C. Funk, Nathan D. Price

**Affiliations:** Center for Bioinformatics, School of Information Science and Engineering, Central South University, Changsha, Hunan Province 410083, P.R. China; Institute for Systems Biology, Seattle, WA 98109, United States

## Abstract

**Summary:** Detecting intron retention (IR) events is emerging as a specialized need for RNA-seq data analysis. Here we present iREAD (**i**ntron **RE**tention **A**nalysis and **D**etector), a tool to detect IR events genome-wide from high-throughput RNA-seq data. The command line interface for iREAD is implemented in Python. iREAD takes as input an existing BAM file, representing the transcriptome, and a text file containing the intron coordinates of a genome. It then 1) counts all reads that overlap intron regions, 2) detects IR vents by analyzing features of reads such as depth and distribution patterns, and 3) outputs a list of retained introns into a tab-delimited text file. The output can be directly used for further exploratory analysis such as differential intron expression and functional enrichment. iREAD provides a new and generic tool to interrogate poly-A enriched transcriptomic data of intron regions.

**Availability:** www.libpls.net/iread

## 1 INTRODUCTION

Historically being considered as transcriptional noise or ‘junk’, intron retention (IR) has recently been shown to carry out important biological functions such as regulating gene expression that is coupled with nonsense mediated decay (Braunschweig, et al., 2014), producing novel isoforms (Bell, et al., 2008) and targeting specific cell compartments (Buckley, et al., 2011). It has thus been gaining recent interest, especially as it relates to a putative role in health and disease. Tumor-specific IRs have been found to be overexpressed in lung adenocarcinoma tumors (Zhang, et al., 2014), and IRs were mostly increased in an analysis across sixteen cancers (Dvinge and Bradley, 2015). IR appears to be a widespread mechanism of tumor-suppressor inactivation based on integrative analysis of exome and RNA sequencing data (Jung, et al., 2015). IR was also found to regulate gene expression in biological processes such as normal granulocyte differentiation (Wong, et al., 2013), CD4+ T cell activation (Ni, et al., 2016), and terminal erythropoiesis (Pimentel, et al., 2016). Functions of IR were systematically discussed in two recent reviews (Ge and Porse, 2014).

Next generation sequencing has resulted in a vast amount of RNA-seq data, which provides a rich resource for the detection of IR in combination with bioinformatics tools. However, to the best of our knowledge, only a few computational tools have been developed thus far—and these tools are either not freely available or have limitations. The IRfinder method and the methods described in (Boutz, et al., 2015) and (Dvinge and Bradley, 2015) are not publicly available. IRcall and IR classifier are also not currently available, possibly due to website changes (Bai, et al., 2015). The recently developed KMA (“Keep Me Around”) is an efficient method for IR detection (Pimentel, et al., 2015). It involves transcript quantification from the command line followed by intron retention analysis in R, which brings some inconveniences for changing work environments. Also, flatly distributed retained intron reads, a common feature of IR, are not well identified in KMA.

In this work, we present iREAD, a simple-to-use command line software for the identification of IR from poly-A enriched RNA-seq data (single-end and paired-end). iREAD takes existing BAM files and a gene annotation file (i.e. ENSEMBL) file as input, count reads in introns and outputs IR events in a tab-delimited file by applying a set of filters (Figure 1).

**Figure 1.**
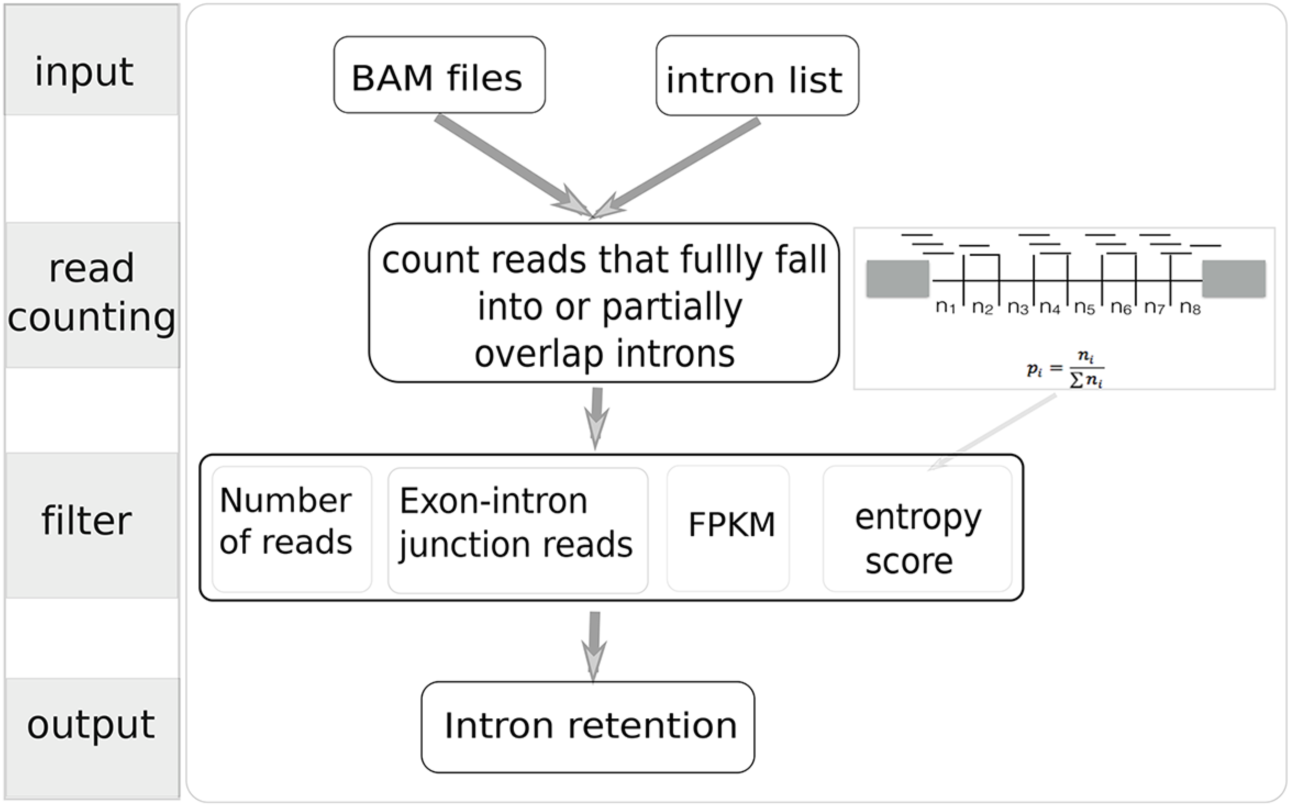
A schematic of iREAD for detecting intron retention. With a BAM file and intron annotation file as input, iREAD first counts intronic reads followed by identifying IR events through a set of filters on read counts, junction reads, FPKM and entropy.

## 2 METHODS

### 2.1 Algorithm

iREAD takes as input an existing BAM file representing the transcriptome and a text file containing the intron coordinates of a genome (Figure 1). Since introns of one splice isoform may overlap with exons of other isoforms, we first seek to identify intron regions that do not overlap with any exons of any isoforms. Such introns are called independent introns for convenience, which are calculated by merging exons of all isoforms and genes of a given genome followed by subtracting them from spanning regions of genes using Bedtools. Using ENSEMBL gene models (GTF format), we identified independent introns together with their coordinates and parent-gene information for humans and mice, and provided them in the iREAD package. Independent introns of any other species can be identified in the same way. Therefore, iREAD only identifies novel IR events not found in the annotation file by default. But known IRs can also be analyzed by manually adding them to the input intron annotation file.

Next, to reduce computational cost, we extract the independent intronic regions from the BAM files using Samtools and count the read depth with Bedops. Since the resulting reads include spliced reads that can span but do not physically overlap introns, we developed a Perl script to count reads that fully reside in or partially overlap with the pre-calculated independent introns by considering both the read-spanning length (from the reference genome) and the coordinates of the independent introns. Specifically, iREAD counts and records the number of exon-intron junction reads, which are valuable for detecting IR. Because reads in retained introns are often flatly distributed across the whole intron region, we use a score to characterize the ‘flatness’ of intronic reads based on information theory. We divide each intron into eight bins, count the number of reads in each bin, and record the number of reads in a vector, (n_1_, n_2_, …, n_8_). This vector is then converted to a probability mass distribution by normalization using the formula 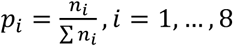. The entropy of this distribution is calculated by: 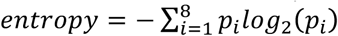

Because the maximal entropy for eight bins is three, we divide the entropy by three and normalize it to be in the range [0,1], called the normalized entropy score (NE-score) for convenience.

We recorded the number of total reads and junction reads, and the NE-score of each intron. FPKM is also calculated. Based on these features, we can define threshold values to filter for IR events. In the iREAD package, high default threshold values (NE-score≥0.9) are used to identify intron retention events conservatively and reliably.

We validated the effectiveness of iREAD using existing IRs annotated in the mouse GTF file (Ensembl version 75). We first removed a total of 12,640 isoforms corresponding to 5,627 genes that are annotated as ‘retained_intron’ in the GTF file, and applied iREAD to a mouse brain sample (https://www.synapse.org/#!Synapse:syn4486837, sample ID:256520). Of note, we found that 1197 (28.1%) out of the 4,266 IRs identified in this single sample were annotated in the GTF file, suggesting that our method was able to detect IRs reliably. At the gene level, these 1,197 IRs fall into 661 genes, covering 11.7% of all 5,627 genes with annotated IR. As the annotation file is a catalog containing gene models for all cell types, IR events may be masked by the existence of tissue-specific isoforms.

As an example of iREAD’s capabilities, a high-confidence retained intron (chr19:5496402-5496985) of *Snx32* (ENSMUSG00000056185) is shown in Figure 2, with an NE-score of 0.995 and coverage by 420 reads of which 157 are junction reads.

**Figure 2.**
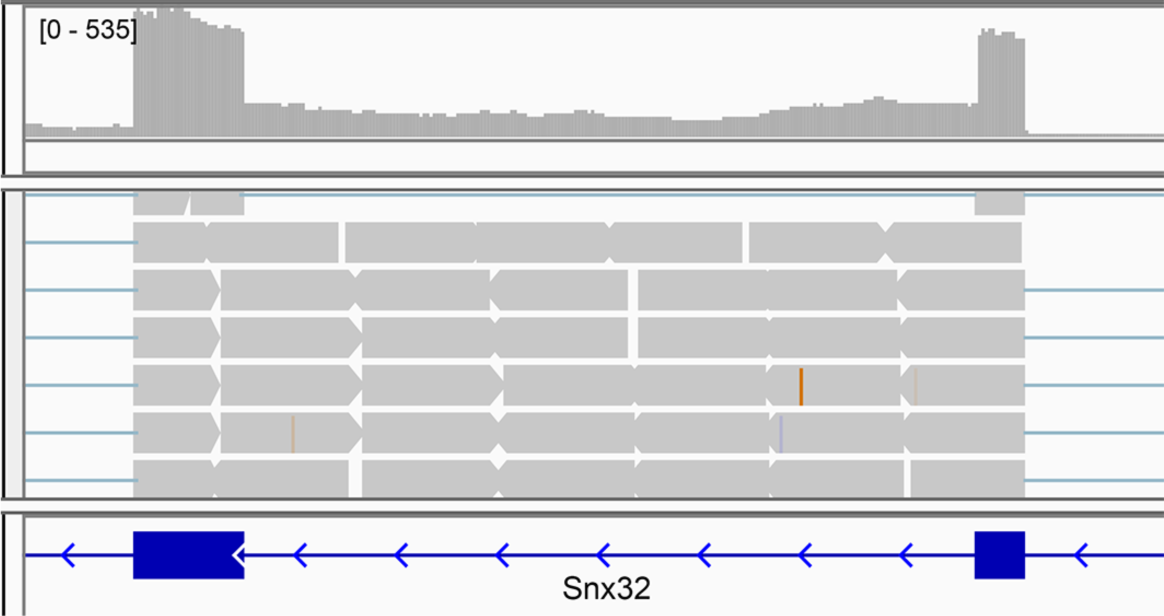
Example of a retained intron (chr19:5496402-5496985) of *Snx32* in a mouse RNA-seq sample visualized in IGV.

### 2.2 The software package

We implemented the iREAD pipeline on the command line using Python. It takes a BAM file and a text file of intron coordinates as input. We implemented multiple options for users to tune parameters of IR filters. Execution of a single command is needed for detecting retained introns from a RNA-seq sample, making it very easy to use. Along with the package, an example is provided for testing. Using the above-mentioned deeply sequenced mouse sample with 133 million reads (9.9G BAM file), it took approximately 30 minutes to identify IR events using a Mac laptop with an Intel Core i7 4-core CPU and 16GB memory. We anticipate that the time could be reduced significantly using routine workstations, with more cores and memory.

## 3 CONCLUSION

By quantifying intronic reads and mining their characteristics, we designed the iREAD algorithm for genome-wide detection of intron retention events from RNA-seq data and implemented it as command line software using Python. It works with both single-end and paired-end sequencing data that has been prepared using poly-A enrichment. The resulting intron retention data can be further explored in various ways such as differential expression and functional enrichment, allowing for its use in many fields including the search for disease-associated gene expression signatures. iREAD provides a new and generic tool to interrogate the previously largely neglected intronic regions from the angle of view of intron retention.

## ACKNOWLEDGEMENTS

Funding: This work was supported by the NIA U01AG006786 (NDP) and start-up funding (NO. 502041004) from Central South University (HDL).

